# Targeted inactivation of spinal α2 adrenoceptors promotes paradoxical anti-nociception

**DOI:** 10.1101/2025.02.06.636935

**Authors:** Javier Lucas-Romero, Maria Fernanda Bandres, Jacob Graves McPherson

## Abstract

Noradrenergic drive from the brainstem to the spinal cord varies in a context-dependent manner to regulate the patterns of sensory and motor transmission that govern perception and action. In sensory networks, it is traditionally assumed that activation of spinal α2 receptors is anti-nociceptive, while spinal α2 blockade is pro-nociceptive. Here, however, we demonstrate *in vivo* in rats that targeted blockade of spinal α2 receptors can promote anti-nociception. The anti-nociceptive effects are not contingent upon supraspinal actions, as they persist below a chronic spinal cord injury and are enhanced by direct spinal application of antagonist. They are also evident throughout sensory-dominant, sensorimotor integrative, and motor-dominant regions of the gray matter, and neither global changes in spinal neural excitability nor off-target activation of spinal α1 adrenoceptors or 5HT_1A_ receptors abolished the anti-nociception. Together, these findings challenge the current understanding of noradrenergic modulation of spinal nociceptive transmission.

## INTRODUCTION

Noradrenergic drive from the brainstem to the spinal cord is a critical regulator of both sensory and motor transmission.(1, 2) Central to this role are α2 adrenergic receptors, the canonical inhibitory adrenergic receptor in the central nervous system.(3) In spinal circuits mediating nociceptive transmission, activation of pre-synaptic α2 heteroceptors on the central terminals of primary afferent neurons reduces glutamate release onto second order sensory neurons, where concurrent activation of post-synaptic α2 heteroceptors results in membrane hyperpolarization.(4–7) This potent inhibitory mechanism has led to the consensus that activation of spinal α2 adrenoceptors is anti-nociceptive, motivating substantial investigation of therapeutics that directly activate spinal α2 receptors (e.g., clonidine) or increase spinal norepinephrine levels (e.g., serotonin-norepinephrine reuptake inhibitors, SNRIs) as analgesics or adjuvants.(8, 9)

In addition to α2 heteroreceptors, α2 adrenoceptors are also located on bulbospinal noradrenergic neurons themselves. Functioning as inhibitory autoreceptors, activation of these α2 adrenoceptors reduces intrinsic discharge rate and spinal norepinephrine release.(10) Curiously, while α2 autoreceptors are well-documented somatodendritically(11–14) and at supraspinal axon terminals,(15) they have never been unequivocally identified at spinal terminals.(16) Indirect evidence supporting their existence in the spinal cord is drawn from spinal slice and synaptosome preparations, in which α2 adrenergic drugs can manipulate norepinephrine release in a manner consistent with autoinhibition.(17–19) However, these findings are challenged both by opposing observations *in vivo*(20) and by a lack of support from direct assays of their existence. For example, immunohistochemical studies indicate that spinal α2 adrenoceptors do not co-localize with neurons containing dopamine β hydroxylase, tyrosine hydroxylase, or phenylethanolamine N-methyltransferase,(14, 21, 22) and neither toxic lesions of noradrenergic fibers nor complete spinal cord transection substantially reduces the number of spinal α2 adrenoceptor binding sites *in vivo.*(23, 24)

In contrast to this understanding of α2 adrenergic actions in the spinal cord, here we demonstrate that targeted blockade of spinal α2 adrenoceptors *writ large* paradoxically depresses nociceptive transmission *in vivo*. The anti-nociceptive effects of α2 adrenergic blockade persisted below a chronic spinal cord injury that damages ascending and descending pathways, and it was evident throughout sensory-dominant, sensorimotor integrative, and motor-dominant regions of the gray matter. These anti-nociceptive effects could not be explained by global changes in spinal neural excitability or by off-target activation of α1 adrenoceptors or 5HT_1A_ receptors, both of which are known to modulate nociception independently of α2 adrenoceptors. Together, these findings motivate a re-evaluation of our understanding of noradrenergic modulation of spinal nociceptive transmission.

## RESULTS

### Systemic blockade of α2 adrenoceptors reduces spinal population responsiveness to nociceptive transmission *in vivo*

We first established the effect(s) of systemic blockade of α2 adrenoceptors on spinal nociceptive transmission (*n*=5 rats). To do so, we quantified the discharge rate of multi-unit intraspinal neural activity in response to noxious mechanical stimulation (painful pinches) of the ipsilateral hindpaw. Multi-unit intraspinal neural activity captures synaptic dynamics at the neural population level, reflecting the many ways that primary afferent feedback is sculpted as it is integrated into segmental networks. We accessed intraspinal neural activity *in vivo* using dual-shank microelectrode arrays (32 channels per array; Neuronexus, Inc.) implanted into the lumbar enlargement at the L5 dorsal root entry zone. To overcome ambiguities associated with prior investigations of spinal α2 adrenoceptor functions, we used a targeted pharmacological probe, RX821002, to block α2 adrenoceptors. RX821002 has ∼100-200x greater affinity for α2_D_ adrenoceptors than α1 adrenoceptors, ∼75-150x greater affinity for α2_D_ adrenoceptors than 5-HT_1A_ receptors (where it is an *antagonist*), and effectively no affinity for imidazoline receptors.(25)

Prior to α2 adrenoceptor blockade, 31 recording electrodes (across rats) exhibited clear pinch-evoked increases in multi-unit discharge rate over that of spontaneous, ongoing neural transmission. We then systemically blocked α2 adrenoceptors via intraperitoneal injection of the antagonist RX821002. Surprisingly, within 2-5 min of RX821002 injection, average pinch-evoked discharge rates decreased in 21/31 electrodes and 4/5 rats (**Fig. 1**). Average pinch-evoked discharge rates increased in the remaining rat, which itself contained a plurality of the 10 electrodes exhibiting increased discharge rates post-drug. As a result of this dispersion, however, the overall main effect of drug on discharge rate was not significant (pre-drug estimated marginal mean: 37.10 ± 8.45; post-drug: 31.93 ± 8.50; *p* = 0.25).

**Figure 1.**
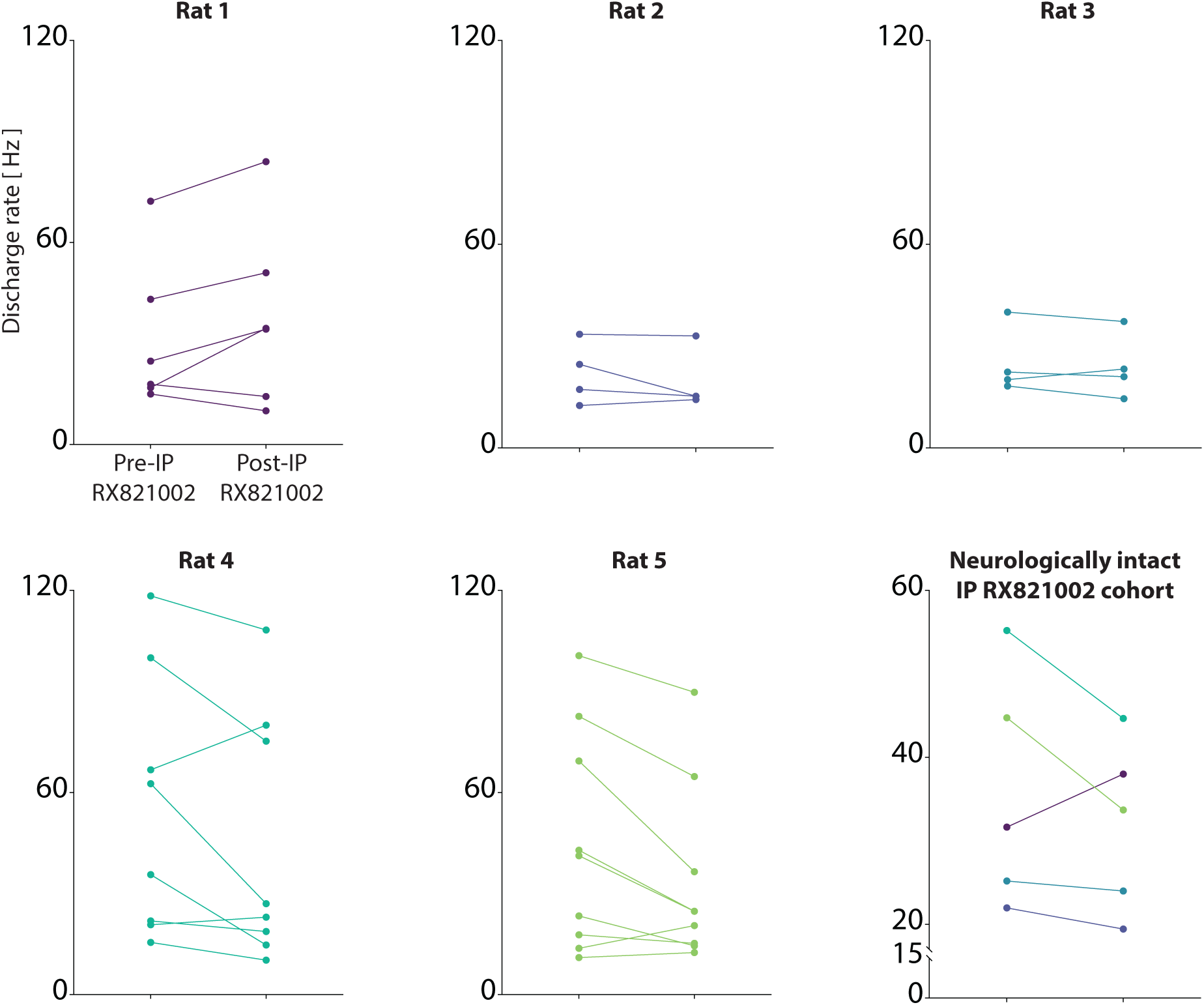
Systemic blockade of α2 adrenoceptors can reduce spinal responsiveness to nociceptive sensory feedback. For all subplots, Y-axis: multi-unit discharge rate; X-axis: (left) prior to intraperitoneal administration of the selective α2 adrenergic blocker, RX821002, (right) following RX821002 administration. For subplots of data from individual rats, each line represents the discharge rate of an electrode determined to be pinch-responsive prior to drug administration. For the subplot of cohort-level data, data are presented rat-level means. Across the cohort – pre-drug estimated marginal mean: 37.10 ± 8.45; post-drug: 31.93 ± 8.50; *p* = 0.25.

### Anti-nociceptive effects of α2 adrenoceptor blockade persist below a chronic spinal cord injury

Because systemic administration of drug in neurologically intact animals could have simultaneously impacted spinal and supraspinal α2 adrenoceptors, we next conducted a series of experiments to determine if *spinal* α2 adrenoceptor inactivation was sufficient to promote anti-nociception. First, we investigated the potential impact of α2 adrenoceptor blockade on nociceptive transmission below a chronic thoracic spinal cord injury (SCI) that results in permanent bilateral hindlimb impairments (*n*=7 rats). This injury reduces norepinephrine release below the lesion in two ways: (1) directly, by damaging descending noradrenergic fibers, and (2) indirectly, by compromising endogenous ascending-descending nociceptive controls. Consequently, we predicted that SCI would reduce or abolish the anti-nociceptive actions observed during systemic administration of RX821002 in neurologically intact animals.

At 6 weeks post-SCI, when spontaneous sensorimotor recovery had stabilized, we conducted a terminal electrophysiological study to determine the effect of α2 adrenoceptor blockade on spinal nociceptive transmission. We again implanted microelectrode arrays into the lumbar enlargement (at the L5 dorsal root entry zone) and characterized pinch-evoked changes in multi-unit discharge rate. Prior to systemic administration of RX821002, pinch-evoked increases in multi-unit discharge rate were detected in 135 recording electrodes across the SCI cohort, with an estimated marginal mean increase of 32.97 ± 6.97 Hz (**Fig. 2**). We then administered RX821002 intraperitoneally, as before, and examined population-level spinal responsiveness to pinch. Contrary to our prediction, RX821002 significantly decreased pinch-evoked discharge rate across animals, despite the damage to ascending and descending fibers (27.81 ± 9.66 Hz; *p* = 0.04).

**Figure 2.**
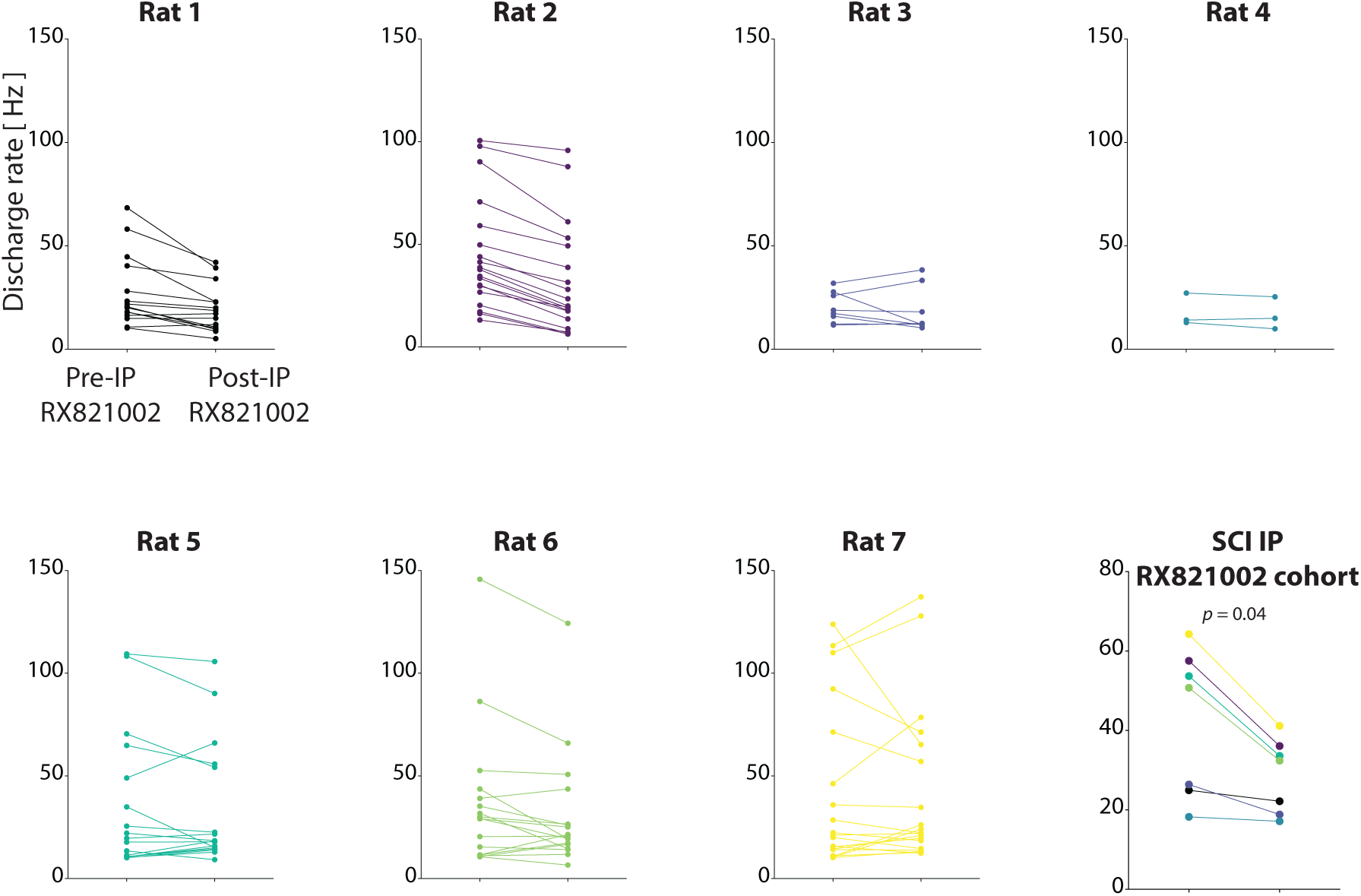
Systemic blockade of α2 adrenoceptors reduces spinal responsiveness to nociceptive sensory feedback below a chronic spinal cord injury. For all subplots, Y-axis: multi-unit discharge rate; X-axis: (left) prior to intraperitoneal administration of the selective α2 adrenergic blocker, RX821002, (right) following RX821002 administration. For subplots of data from individual rats, each line represents the discharge rate of an electrode determined to be pinch-responsive prior to drug administration. For the subplot of cohort-level data, data are presented rat-level means. Across the cohort – pre-drug estimated marginal mean: 32.97 ± 6.97 Hz; post-drug: 27.81 ± 9.66 Hz; *p* = 0.04.

### Anti-nociceptive potency is enhanced by direct spinal α2 adrenoceptor blockade

It was difficult to reconcile why the anti-nociceptive efficacy of α2 adrenoceptor blockade was comparable between neurologically intact animals and those with chronic SCI. Indeed, the effective modulatory capacity of noradrenergic fibers spared by SCI is presumably lower than that of fully intact tracts. However, there is evidence that spinal α2-mediated antinociception is driven by descending fibers coursing through the ventral funiculus and not the dorsal funiculus.(26, 27) Given that we utilized a dorsal spinal contusion model, we reasoned that one explanation for the persistence of anti-nociception below the lesion could be differential sparing of ventral vs. dorsal noradrenergic fibers.

To mitigate this confound, as well as that of potential supraspinal actions, we next used a drug-embedded packet to deliver RX821002 directly to the dorsal surface of the spinal cord, immediately adjacent to the microelectrode array implantation site. By incising and reflecting the spinal meninges prior to application of the packet, this approach prevented cerebrospinal fluid from shunting the drug to the ventral aspect of the spinal cord, as could happen with standard intrathecal drug administration. And because drug delivery was confined to the spinal cord, this approach prevented binding to the supraspinal α2 autoreceptors known to promote top-down inhibition.(28)

We tested the impact of direct spinal α2 adrenoceptor blockade on nociceptive transmission in another cohort of neurologically intact rats (*n* = 5). We reasoned that using neurologically intact rats would prevent additional confounds due to variability in the proportion of spared descending noradrenergic fibers between animals with SCI. By extension, we also reasoned that it would more directly decouple spinally mediated effects – whether anti- or pro-nociceptive – from supraspinally mediated anti-nociception.

Prior to drug administration, clear pinch-evoked increases in discharge rate were evident in 30 electrodes. The estimated marginal mean increase in discharge rate during these events was 30.80 ± 6.01 Hz (**Fig. 3**). We then applied RX821002 (30µg/10µl) and repeated the experimental sequence. Remarkably, direct spinal application of RX821002 also depressed nociceptive transmission (estimated marginal mean increase of 17.33 ± 6.00 Hz; *p* < 0.0001), and to a greater extent (44%) than that realized with systemic α2 adrenoceptor blockade, which plausibly could have engaged supraspinal α2 autoreceptors known promote anti-nociception.

**Figure 3.**
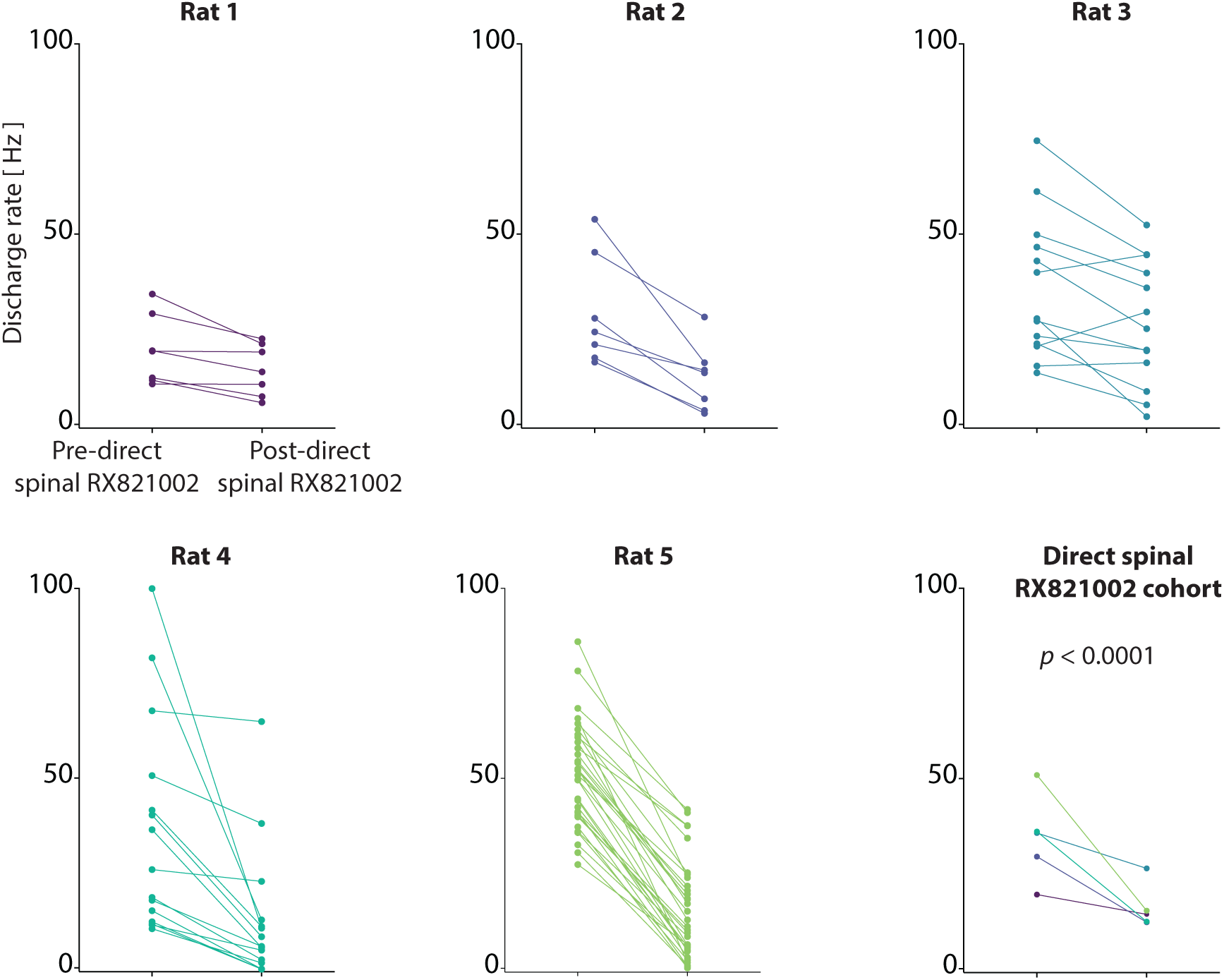
The anti-nociceptive potency of targeted α2 adrenergic blockade is enhanced by direct spinal application of the antagonist RX821002. For all subplots, Y-axis: multi-unit discharge rate; X-axis: (left) prior to direct spinal application of the selective α2 adrenergic blocker, RX821002, (right) following direct spinal RX821002 administration. For subplots of data from individual rats, each line represents the discharge rate of an electrode determined to be pinch-responsive prior to drug administration. For the subplot of cohort-level data, data are presented rat-level means. Across the cohort – pre-drug estimated marginal mean: 30.80 ± 6.01 Hz; post-drug: 17.33 ± 6.00 Hz; *p* < 0.0001.

To ensure that the anti-nociceptive effect of α2 adrenoceptor blockade was indeed driven by drug and not spuriously by the presence of the packet on the spinal cord itself, we conducted two additional sets of control experiments. First, we verified that the effect of drug was reversible. To do so, in 3 additional rats we characterized the pharmacokinetic profile of RX821002-mediated anti-nociception. We delivered pinches (as above) at 30-s intervals for 1 hr and quantified the timecourse over which nociceptive transmission was depressed by direct spinal application of RX821002 (**Fig. 4A**). After 15 minutes of pre-drug baseline pinches, the drug embedded packet was placed on the dorsal surface of the exposed spinal cord. Clear decreases in responsiveness to pinch became evident within the first ∼2-5 min after drug application, reaching a maximum effect within ∼5-10 min. The anti-nociceptive effect then stabilized before washing out over the ensuing ∼30 min. At 40 min after drug administration, the anti-nociceptive effect was nearly absent, suggesting that all drug had been metabolized and the effect was not due to the physical presence of the packet.

**Figure 4.**
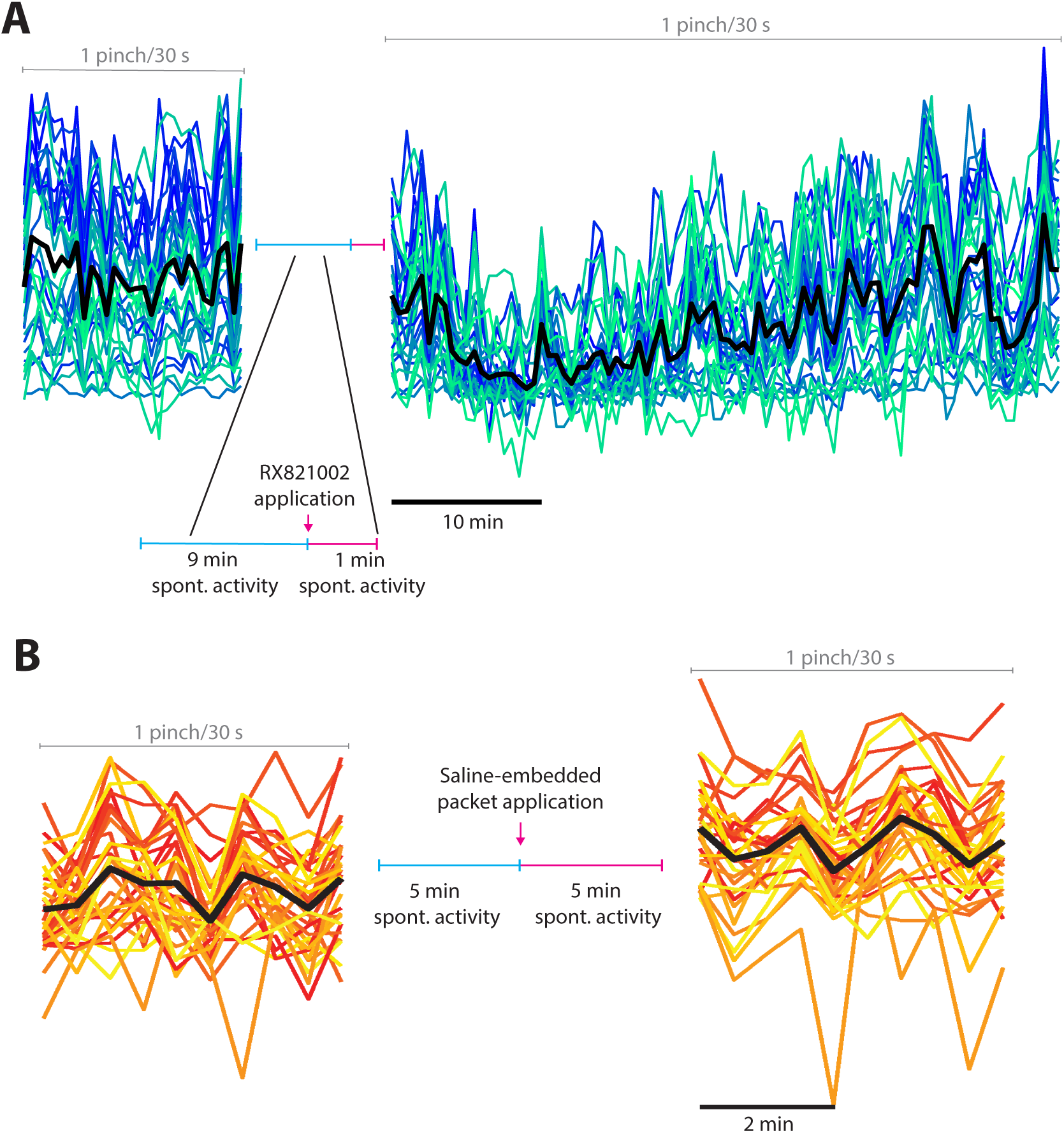
The anti-nociceptive actions of direct spinal RX821002 are reversible and not attributable to the method of drug administration. (A) Multi-unit discharge rate in response to pinch and direct spinal RX821002 administration across all channels of the MEA in one representative rat. Each line color depicts a discrete electrode channel, with the bold black line depicting the channel-level mean discharge rate. The vertical scale is the same for both pre- and post-drug responses, and the horizontal scale likewise applies to both trial segments. (B) Multi-unit discharge rate in response to pinch and direct spinal administration of a saline-embedded packed across all channels of the MEA in one representative rat. Each line color depicts a discrete electrode channel, with the bold black line depicting the channel-level mean discharge rate. The vertical scale is the same for both pre- and post-drug responses, and the horizontal scale likewise applies to both trial segments.

Next, we repeated the procedure using a saline-embedded packet instead of RX821002 (*n* = 7 additional rats; **Fig. 4B**). In contrast to the findings with RX821002, we found no differences in responsiveness to nociceptive sensory feedback after application of the saline packet (pre-saline discharge rate: 32.39 ± 4.16 Hz; during saline: 30.80 ± 4.16 Hz; *p* = 0.16), thus confirming that the effect was unique to drug and not the experimental preparation.

### Single-unit nociceptive responsiveness is also diminished by spinal α2 adrenoceptor blockade

Next, we sought to determine whether the observed population-level anti-nociceptive effect of spinal α2 adrenoceptor blockade masked an enhanced responsiveness specifically in discrete nociceptive-specific and wide dynamic range neurons. To this end, we used computational approaches to discriminate single-unit neural activity from the electrodes that exhibited population-level responsiveness to nociceptive feedback (*n*=5 rats). Across electrodes, we identified 111 well-isolated nociceptive-specific and wide dynamic range neurons that could be reliably tracked from pre- to post-drug administration. For each, we created spike count histograms (0.5 ms bin width) of pinch-evoked spike discharge events and extracted the peak discharge count per pinch event. We then computed the average peak spike count over the 5 min immediately prior to drug application and from 5-10 min after drug application (**Fig. 5A**). Across neurons, the average peak pre-RX821002 count was 14.75 (±13.14) per 0.5 ms. Between 5-10 min after drug application, peak discharge count decreased to 8.62 (±9.50)/0.5 ms, significantly lower than prior to drug application (*p*<0.0001). The depressant effect of RX821002 on single unit nociceptive responsiveness was nearly ubiquitous, with only 5 of 111 neurons (4.5%) exhibiting increased nociceptive responsiveness following drug administration.

**Figure 5.**
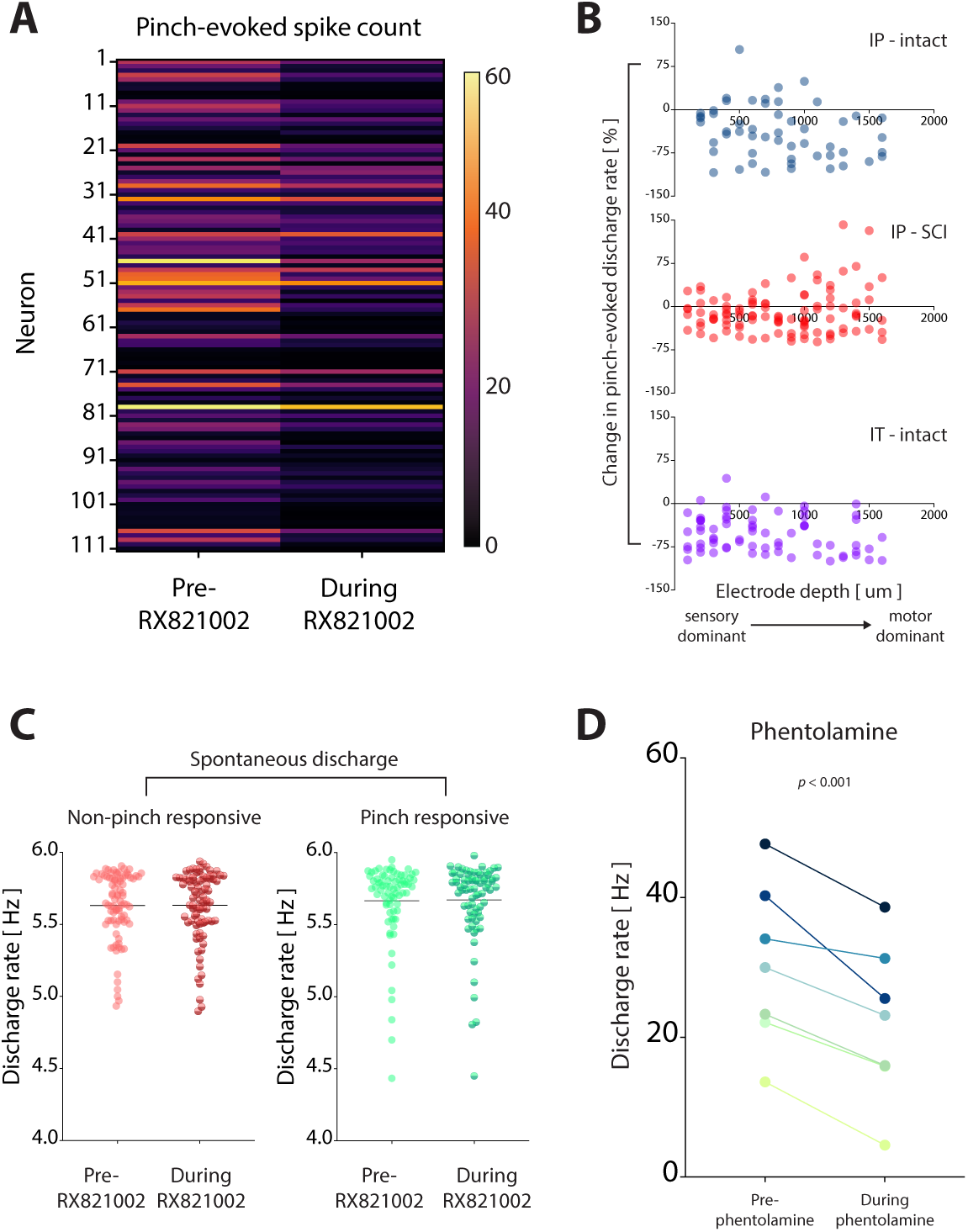
The anti-nociceptive effects of spinal α2 adrenergic blockade neural span scales and functional regions, are not accompanied by changes in global neural excitability, and persist despite concurrent α1 adrenergic blockade. (A) Single-unit intraspinal responses to nociceptive sensory feedback. Y-axis: neuron number. Each line represents a discrete, well-isolated neuron extracted from multi-unit patterns of neural transmission. X-axis: (left) prior to RX821002 administration; (right) following RX821002 administration. Colormap indicates the mean number of spikes per neuron per pinch. Warmer colors indicate more spikes, i.e., greater responsiveness, and darker colors represent fewer spikes, i.e., less responsiveness. *N* = 111 neurons from *N* = 5 rats. (B) RX821002-induced changes in pinch-evoked discharge do not systematically vary across region of the L5 gray matter for any of the cohorts tested. Y-axis for all subplots: percent change in multi-unit discharge rate following RX821002 administration; X-axis: depth of electrode in the spinal gray matter. (C) The discharge rate of spontaneous neural transmission does not change in either non-pinch responsive (left) or pinch-responsive (right) neurons following administration of RX821002. Y-axis: multi-unit discharge rate; X-axis: pre- or during drug. (D) Direct spinal administration of the α1/α2 adrenergic antagonist does not reverse the anti-nociceptive effect of α2 adrenergic blockade alone. Y-axis: multi-unit discharge rate; X-axis: (left) prior to direct spinal administration of phentolamine; (right) following phentolamine administration. Data are presented rat-level means. Estimated marginal means: 30.51 ± 4.51 pre-drug vs. 22.31 ± 4.51 post-drug; *p* ≪ 0.001.

### Inhibitory effect of spinal α2 adrenoceptor blockade is not relegated to overt spinal pain processing networks

The spinal distribution of α2 adrenoceptors(29) and descending noradrenergic fibers(30) is not limited to primary afferents and second order sensory neurons in the dorsal horn, nor are the segmental pathways that support nocifensive behaviors. Both span sensory-dominant, sensorimotor integrative, and motor-dominant regions of the spinal gray matter. Thus, we next asked whether the observed anti-nociceptive effects of α2 adrenoceptor blockade could likewise be observed throughout the gray matter of a given spinal segment.

We first identified the regions of the gray matter in which decreased pinch-evoked discharge rates became evident following α2 adrenoceptor blockade. Because we configured the microelectrode arrays to span the full dorso-ventral and medio-lateral extent of a hemi-segment of the spinal cord, we were able to simultaneously characterize this regional distribution in each animal. In all animals, regardless of whether neurologically intact or with chronic SCI, and regardless of the route of drug administration, we found that decreased pinch-evoked discharge rates extended from the most superficial recording sites in the dorsal horn to the deepest recording sites in the ventral horn, located amongst the spinal motor pools.

This finding indicated that α2 adrenoceptor blockade influenced multiple functional and structural networks within the spinal cord. However, left unresolved was the question of whether the depressant effect was uniform across regions. To address that question, we regressed the magnitude of change in pinch-evoked discharge against the dorso-ventral depth at which the change was observed. Although it was not clear *a priori* whether systematic regional differences would be evident, it was reasonable to speculate that the depressant effect might be most pronounced in the dorsal horn, where α2 adrenoceptors are most densely located. However, we found no dependency of the magnitude of depression on electrode depth for any cohort (**Fig. 5B**), suggesting that spinal α2 adrenoceptor blockade exerted a surprisingly isotropic pattern of population-level anti-nociception.

### Anti-nociceptive effect of spinal α2 adrenoceptor blockade is not driven by a global change in spinal excitability

There is ambiguity as to whether the spinal cord receives a tonic noradrenergic drive during anesthesia-induced unconsciousness.(25) If so, then either systemic or direct spinal RX821002 could have indiscriminately altered the backdrop of spinal excitability and confounded the interpretation of α2 blockade as anti-nociceptive. Conversely, in the absence of a tonic noradrenergic drive, α2 adrenoceptor blockade would not be predicted to alter overall spinal excitability given that there are no endogenous spinal sources of norepinephrine.

To understand whether a broad, drug-induced shift in spinal excitability could have contributed to the observed anti-nociception, we focused instead on spontaneous neural transmission. We quantified spontaneous population-level discharge rates in electrodes not responsive to nociceptive sensory feedback and, separately, those in which clear pinch-evoked responses had been evident (**Fig. 5C**). Prior to direct spinal application of RX821002, the spontaneous discharge rate of non-responsive channels was 5.61 ± 0.18 Hz. This rate remained statistically invariant to α2 adrenoceptor blockade, averaging 5.61 ± 0.18 Hz from 5-10 min post-RX821002 application (*p* = 0.92). The spontaneous discharge rate of pinch-responsive electrodes was likewise stable after drug application (5.67 ± 0.12 Hz vs. 5.68 ± 0.12 Hz; *p* = 0.46), together suggesting that the anti-nociceptive effect of α2 adrenoceptor blockade was not the biproduct of a broad, drug-induced change in spinal neural excitability,

### Opportunistic activation of α1 adrenoceptors does not explain the anti-nociceptive effects of spinal α2 adrenoceptor blockade

Although α1 adrenoceptors are biophysically excitatory, they contribute to spinal anti-nociception via their location on inhibitory glycinergic and GABA-ergic neurons in the superficial dorsal horn.(31) Thus, it was necessary to understand whether free norepinephrine binding to spinal α1 adrenoceptors could have mediated the anti-nociceptive effects observed when α2 adrenoceptors were occupied by drug. To test this hypothesis, we quantified the effect of phentolamine, a nonspecific α1/α2 adrenergic antagonist, on spinal responsiveness to nociceptive sensory feedback. In 7 additional neurologically intact animals, we administered phentolamine directly to the spinal cord via drug-embedded packet, recording multi-unit neural activity prior to and following drug administration. Despite the concurrent inactivation of α1 and α2 adrenoceptors, the net effect of phentolamine in all animals remained anti-nociceptive (**Fig. 5D**; significant main effect of drug on multi-unit responsiveness; estimated marginal means: 30.51 ± 4.51 pre-drug vs. 22.31 ± 4.51 post-drug; *p* ≪ 0.001). This finding suggests that opportunistic binding of norepinephrine to α1 adrenoceptors did not cause the anti-nociceptive effects observed during α2 adrenoceptor blockade.

## DISCUSSION

Our primary finding is that targeted blockade of spinal α2 adrenoceptors can reduce spinal nociceptive transmission *in vivo*. These actions appear to be mediated by α2 adrenoceptors in the spinal cord, not the brainstem, both because the anti-nociceptive effect persisted below a chronic spinal cord injury and because its potency was enhanced when drug was delivered directly to the spinal cord, bypassing supraspinal nuclei. We also found that the depressive effect of α2 adrenoceptor blockade was robustly expressed throughout multiple structural and functional spinal networks involved in nociception and could not be explained by global changes in spinal excitability or by activation of spinal α1 adrenoceptors or 5HT_1A_ receptors.

Given the relative consistency of the anti-nociceptive effect observed here, it is surprising that direct anti-nociceptive effects of spinal α2 adrenergic blockade have not previously been reported – particularly in studies utilizing RX821002 or the related compound atipamezole. However, several experimental design choices could have contributed. The first is scale. Prior studies have focused either on the excitability of primary afferents and second order sensory neurons in the superficial dorsal horn or on movement-related components of nocifensive reflexes.(5–7, 17–19, 25) We instead focused on neural population-level patterns of spinal nociceptive transmission. At the most reductionist scale, it is difficult to dispute that *activation* of spinal α2 receptors is inhibitory. Indeed, they are G_I/o_ protein-coupled receptors, and activation of pre-synaptic α2 receptors on the terminals of Aδ and C fiber primary afferents reduces glutamate release onto cell bodies in the substantia gelatinosa.(5, 7) In tandem, activation of post-synaptic α2 receptors on second order neurons hyperpolarizes their resting membrane potential,(6) further reducing nociceptive transmission.

However, our results suggest that spinal population-level nociceptive transmission is not wholly dictated by synaptic interactions at this first synapse, and in cases may be in opposition. In that regard, our findings are philosophically consistent with the recent observation that global activation of low threshold Aβ mechanoreceptors is anti-nociceptive whereas their local activation is pro-nociceptive.(32) They are likewise consistent with the observation that population-level activity, but not single-cell activity, encodes noxious stimuli in spinothalamic tract neurons(33) and the anterior cingulate cortex.(34) In this context, it was somewhat surprising that we found congruency between the responses of individual nociceptive specific and wide dynamic range neurons and population-level neural transmission. Presumably, however, this finding reflects the *in-situ* nature of our recordings, in which even single-unit spiking reflects the integrative actions of multiple convergent synaptic inputs.

At the scale of behavior, norepinephrine and α2 adrenergic drugs are also potent modulators of vigilance. Through their supraspinal actions, α2 agonists promote sedation while antagonists promote alertness. Thus, previous reports of α2 adrenergic agonists promoting anti-nociceptive behaviors may have actually arisen from α2-mediated sedation, which reportedly occurs at lower drug doses than anti-nociception.(35) Norepinephrine and α2 adrenergic drugs also have direct effects on motoneuron excitability,(36–39) a particular confound for studies utilizing nocifensive motor reflexes as outcomes. Unexpectedly, we found that α2 adrenoceptor blockade depressed nociceptive responsiveness in motor-dominant regions of the gray matter. It is likely that the inhibited cells were premotor interneurons and not motoneurons, however, given that the animals were deeply anesthetized and lacked clear withdrawal reflexes. Nevertheless, this finding is a notable departure from the pro-nociceptive changes in withdrawal reflexes reported in rabbits following intrathecal administration of RX821002.(25)

Another potential contributor to the divergence between our results and those reported previously is the still prevalent use of non-specific pharmacological probes. Especially problematic is the noradrenergic antagonist yohimbine, which has poor α1/α2 separability, limited ability to differentiate between α2 adrenoceptor subtypes, and a characteristically low affinity for α2_D_ adrenoceptors – the dominant spinal α2 adrenoceptor subtype in mice and rats.(25, 40–42) Yohimbine is also an agonist at the spinal 5-HT_1A_ receptor, activation of which inhibits nociceptive transmission and enhances motor excitability independently of the noradrenergically driven cascade.(43, 44) Thus, the off-target effects of yohimbine may have masked the actions of the very α2 adrenoceptors it has been intended to study.

An additional consideration is the modality of nociceptive transmission under study. Here, we focused on mechanical nociception. In contrast, prior studies have focused primarily on thermal, chemical, and electrically induced nociception.(45–49) Under the nominal assertion that transmission of different sensory modalities involves peripheral nerves and spinal networks that do not uniformly overlap,(50, 51) then at least some of the differential effects of α2 adrenoceptor blockade observed across studies presumably reflects these differences in structural and functional network connectivity, resulting in the emergence of distinct population-level patterns of neural transmission.

At a mechanistic level, the observed anti-nociceptive effects remain enigmatic. Even if α2 autoreceptors do exist in the spinal cord at functionally relevant levels, we have no reason to suspect that RX821002 or phentolamine would preferentially bind to them vs. the α2 heteroreceptors on primary afferents and second order sensory neurons. Indeed, both drugs are relatively non-specific for α2 subtypes (although note the aforementioned preference of RX821002 for α2_D_).(25) Thus, an anti-nociceptive effect driven by this mechanism seems unlikely. An anti-nociceptive effect mediated by 5HT_1A_ receptors(34, 52–55) is also an unlikely candidate, as RX821002 is a pure antagonist at these receptors and appears to be selective for α2 adrenoceptors below 100 μg.(44, 49) Anti-nociceptive actions at α1 adrenoceptors are likewise not implicated, as phentolamine, which inactivates both α1 and α2 adrenoceptors, did not abolish the anti-nociceptive effects observed during α2 blockade alone. Finally, it also seems unlikely that free spinal norepinephrine would have substantially displaced RX821002-bound α2 heteroreceptors, as the affinity of RX821002 for α2 adrenoceptors is considerably greater than that of their endogenous ligand (Ki of 8.1-9.2 for RX821002 vs. 3.6-7.4 for norepinephrine).(56) Thus, absent any as-yet unrecognized actions of RX821002 at non-adrenergic binding sites, it seems reasonable to speculate that the observed anti-nociceptive effects are related in some way to increased release of spinal norepinephrine secondary to blockade of inhibitory α2 autoreceptors.

But what downstream mechanism(s) could have subsequently been engaged? Arguably the simplest explanation would be a difference in the number of antagonist-bound spinal α2 autoreceptors relative to α2 heteroreceptors. The inability to unequivocally identify α2 autoreceptors at the spinal terminals of noradrenergic neurons suggests that they are sparse. If so, it is possible that the drugs occupied a greater proportion of α2 autoreceptors than heteroceptors, leaving available a sufficient number of heteroreceptors for the enhanced free norepinephrine to promote anti-nociception. This dual effect – increased release of norepinephrine coupled with an incomplete α2 heteroreceptor blockade – would be consistent with the observation that low doses of α2 antagonists can enhance norepinephrine, clonidine, and morphine-induced analgesia.(57, 58) However, it should be noted that those studies used antagonist doses considerably lower than those used here, specifically to avoid robust α2 blockade.

Other mechanisms are also possible. Norepinephrine and α2 adrenergic drugs modulate cholinergic interneurons, which can indirectly promote anti-nociception via nitric oxide.(59, 60) It has been suggested that this mechanism is more potent than autoinhibition in typical physiologic circumstances,(18) and if so, then blockade of putative spinal α2 autoreceptors could have augmented anti-nociception through this pathway. There is also evidence that norepinephrine released from post-ganglionic sympathetic nerves excites nociceptive primary afferents in the dorsal root ganglia by activating somatic α2 adrenoceptors;(61) signaling through these afferents can be reduced via α2 adrenergic antagonists.(62, 63) Although to date this effect has only been documented following peripheral nerve injury, the possibility cannot be excluded that the anti-nociceptive effect observed in the present study is also related to this mechanism.

Ultimately, future studies will be required to define the mechanisms by which spinal α2 adrenergic reduced nociceptive transmission. Nevertheless, the present findings challenge the traditional understanding of the functions of spinal α2 adrenergic receptors.

## METHODS

### Animals

Adult male Sprague-Dawley rats were used in this study (*n* = 27). Therefore, the extent to which the findings are relevant to both sexes is unknown. Rats were distributed as follows: experiments involving intraperitoneal injection of RX821002 in neurologically intact rats: 5 rats; spinal cord injury experiments: 7 rats; experiments involving direct spinal application of RX821002: 5 neurologically intact rats; experiments to quantify the pharmacokinetic profile of RX821002: 3 neurologically intact rats; control experiments with saline-embedded spinal packets: 7 neurologically intact rats, of which 6 were also used in phentolamine experiments; and experiments using direct spinal application of phentolamine utilized: 7 neurologically intact rats. All animals were group housed (2-3/cage) with standard food and water available *ad libitum*.

### Study approval

All procedures were approved by the Institutional Animal Care and Usage Committee of Washington University in St. Louis.

### Spinal cord injury

All SCI-related surgical procedures were conducted in an aseptic environment. Heart rate, blood pressure, SpO_2_, and core temperature were monitored from anesthesia induction until animals were transferred to recovery housing following conclusion of the surgical procedures (∼1-1.5 hours) (Kent Scientific, Inc.). Anesthesia was induced with inhaled isoflurane (∼3-4% O_2_, flow rate: 1-2 L/min) and maintained with intraperitoneal injection of ketamine (80 mg/Kg) and xylazine (12 mg/Kg) (boosts as necessary; ketamine at 40 mg/Kg and xylazine at 6 mg/Kg). Once anesthetized, a ∼5 cm midline incision was made in the skin over the dorsal aspect of the vertebral column, centered at the 8^th^ thoracic vertebra. Subcutaneous tissue and muscle were reflected, and musculotendonous attachments to the 8^th^ and 9^th^ thoracic vertebrae (T8/T9) were removed. The caudal half of the T8 lamina and the rostral half of the T9 lamina were removed, revealing the dorsal surface of the spinal cord. With the meninges intact, a spinal cord impactor generated a midline contusion injury at the T8/T9 border (Infinite Horizons Impactor, IH-04000; Precision Systems and Instrumentation, LLC; 200 Kilodynes force and 0 sec of dwell time; 2.5 mm diameter probe located ∼ 3mm from the dorsal cord surface).

Following the contusion, muscle was closed in layers and the skin was closed with suture and surgical staples. Animals were then administered buprenorphine SR (1mg/Kg), warmed lactated Ringer’s solution (3-5mL), and antibiotic (Enrofloxacin 0.5mg/Kg). Animals were then wrapped in sterile cloth and transferred to warmed recovery housing until they awoke from anesthesia. Post-operative care involved daily administration of nutritional supplements, electrolyte replenishers (Bio-Serv), water with antibiotic (Enrofloxacin 0.5ml/L) and sweetener, and warmed lactated Ringer’s solution (3-5mL). The bladder was manually expressed at least 3 times per day until spontaneous voiding reflexes returned.

### Microelectrode array implantation

All animals underwent the terminal procedures described below. In animals with SCI, these procedures occurred 6 weeks post-injury, when spontaneous recovery of sensory and motor function had ceased.

Anesthesia was induced via inhaled isoflurane (∼1-3% O_2_, flow rate: 1-2 L/min) and subsequently maintained with intraperitoneal injection(s) of urethane (1.2 g/kg i.p.). Urethane was chosen due to its ability to preserve the excitability of spinal nociceptive and sensorimotor pathways.(64–66) Vital signs were monitored continuously during the experiment and core temperature was maintained using a feedback-controlled thermal pad (Kent Scientific, Inc.). Rats were given subcutaneous injections of lactated Ringer’s solution every 2 hrs to prevent dehydration.

Under deep, surgical plane anesthesia, a ∼5cm incision was made in the skin over the vertebral column. Muscle and subcutaneous tissue were dissected in layers, and musculotendonous attachments were cleared from the dorsal processes of the T13-L2 vertebrae. The T12 and L3 vertebrate were clamped with locking forceps attached to a custom fixation frame, which was then slightly elevated to prevent respiration cycles from causing vertical displacements of the spinal cord. A dorsal laminectomy was then performed on the T13-L2 vertebral segments and the exposed dura mater incised rostro-caudally and reflected.

The animals were placed on an anti-vibration air table enclosed by a Faraday cage for microelectrode array (MEA) implantation and all subsequent electrophysiological and pharmacological experiments. MEAs consisted of two parallel shanks, each containing 16 discrete, vertically aligned electrodes (electrode area: 177 μm^2^; inter-electrode spacing: 100 μm; NeuroNexus Inc., A2×16). The tips of both shanks were sharpened and the MEAs were custom electrodeposited with activated platinum-iridium (impedance: 4-10 KΩ; Platinum Group Coatings, Inc.). MEAs were also coated with 1,1’-Dioctadecyl-3,3,3’,3’-tetramethylindocarbocyanine perchlorate (Sigma-Aldrich, Inc.) to aid postmortem histological localization.

Each rat received a single MEA implant. The MEA was positioned perpendicular and slightly lateral to the midline at the level of the L5 spinal nerve dorsal root entry zone. Using a custom, multi-axis micromanipulator (Siskiyou, Inc.), the MEA was lowered until the bottom-most electrodes contacted the dorsal surface of the spinal cord, amongst the dorsal roots. We then mapped the L4, L5, and L6 dermatomes by mechanically probing the ipsilateral hindpaw, monitoring dorsal root potentials in real-time. If dorsal root potentials matched receptive fields on the glabrous skin of the ipsilateral hindpaw, corresponding to the L5 dermatome, MEA implantation commenced. If dorsal root potentials were either absent or correlated with another dermatome, the MEA was repositioned, and the mapping procedure was repeated.

After establishing the implant location, the implantation procedure began. The MEA was advanced into the spinal cord in ∼25-50 μm increments, pausing to minimize shear and planar stress on the neural tissue. At a depth of ∼400-500 relative to the dorsal surface of the spinal cord, corresponding to the deep dorsal horn, implantation was halted and the L5 dermatome was re-mapped. If intraspinal neural activity continued to be evoked in response to mechanical stimulation of the desired receptive field, implantation continued. If not, the MEA was withdrawn, and a new implantation site was established. When fully implanted, the deepest electrodes on the MEA were ∼1,600 – 2,000 μm deep to the dorsal surface of the spinal cord, corresponding to the ventral horn, while the dorsal-most electrodes were ∼100-200 μm deep to the surface, residing within the superficial dorsal horn. This configuration enabled simultaneous recording throughout the gray matter of one half of the L5 spinal segment, spanning sensory-dominant, sensorimotor integrative, and motor-dominant regions. After full implantation, the MEA was not removed until all procedures were complete, and the rat was humanely euthanized with an overdose of sodium pentobarbital.

### Data acquisition and trial structure

Trials were designed to enable characterization of the effect(s) of pharmacological challenge on spinal responsiveness to natural nociceptive sensory feedback. Trial structure was the same for all animal cohorts and drugs. Each trial consisted of the following components, in order: (1) 5 min of recording from the MEA without induced sensory feedback (spontaneous neural transmission), (2) ≥ 5 min of pre-drug baseline MEA recording during which nociceptive sensory feedback was induced episodically, (3) drug administration, and (4) 20-30 min of MEA recording during episodic nociceptive feedback. For pharmacokinetic experiments, recording continued for at least 60 min post-drug administration. All phases were contiguous, and neural data was acquired simultaneously from all electrode channels (Ripple Neuro, Inc).

Nociceptive sensory feedback consisted of a series of pinches of the hindpaw ipsilateral to the MEA. Pinches (1-2 s duration each) were delivered once every ∼30 s to avoid tissue damage and central sensitization. Pinch force magnitude was sufficient to induce a robust withdrawal reflex in an unanesthetized rat, and all pinches were delivered by the same experimentalist across cohorts.

### Pharmacology

This study used the noradrenergic α2 antagonist RX821002 (Tocris Biosciences) and the noradrenergic α1/α2 antagonist phentolamine (Sigma). Intraperitoneal injections of RX821002 were administered at 1 g/Kg for both the neurologically intact and SCI cohorts. For direct spinal application of drugs, Kimwipe pledgets were impregnated with either RX821002 (30 µg in 10 µl of Ringer’s solution) or phentolamine (30 μg in 20 μl of Ringer’s solution) and placed directly on the exposed spinal cord immediately adjacent to the MEA. Phentolamine was only delivered via the direct spinal route.

### Multiunit analysis

For each trial, broadband neural activity (μV) was recorded at 30 KHz from all electrode channels of the MEA and stored for offline analysis. Raw data were high pass filtered at 750 Hz using a 4^th^ order, zero-phase Butterworth filter (Matlab; the Mathworks, Inc.). The standard deviation (μV) of each electrode channel was then extracted from the 5 min of spontaneous activity that began each trial. For trial segments that included induced nociceptive transmission, we used a threshold of 3 standard deviations (per channel) above the mean voltage to differentiate potentially pinch-responsive activity from ongoing, spontaneous neural transmission. Suprathreshold activity was then used to compute multi-unit discharge rate, defined as the number of peaks above threshold per 2 s bin. The multi-unit response to pinch was taken as the difference in discharge rate between the 2 s immediately prior to a pinch and the 2 s during the pinch itself. For an electrode to be considered as responsive to pinch, its multi-unit discharge rate was required to increase by a mean of ≥ 10 Hz across pinches prior to drug administration.

### Single-unit analysis

Raw, broadband neural data were pre-processed to remove electrical noise as well as physiological and non-physiological artifacts (e.g., EKG and vibrations, respectively). Cleaned data were then decomposed into spike trains of individual neurons using the wavelet-based spike sorting algorithm, “wave_clus,”(64, 65, 67, 68) implemented programmatically in Matlab. The algorithm was configured as follows: bandpass filter: 1 Hz to 15 kHz; minimum detection threshold: 4 standard deviations [SD] from mean; maximum detection threshold: 25 SD; detection thresholds on both positive and negative deviations; filter order for detection: 4; filter order for sorting: 2.

The resulting spike trains were then analyzed manually to remove any decomposition errors (e.g., predominance of interspike intervals < 2ms, non-physiological action potential shape, inappropriate action potential duration). Neurons not passing the manual verification stage were discarded (typically 5-20 total neurons for a given animal across the 32 channel MEA).

### Experimental design and statistical analysis

As initially conceived, this study was intended to verify that systemic administration of the selective noradrenergic α2 antagonist RX821002 would *increase* spinal responsiveness to nociceptive sensory feedback. Upon finding that this was not uniformly the case (in neurologically intact rats administered intraperitoneal RX821002), we designed the subsequent experiments to further probe this unexpected action of α2 adrenergic blockade.

Estimating the required sample sizes for the follow-on cohorts was complicated by several factors. The most formidable challenge was the lack of comparable data in the literature, both in terms of the observed effect (anti-nociceptive effect of α2 adrenergic blockade rather than a pro-nociceptive effect) and the outcome measures/experimental techniques used. The lack of referent data was further complicated by the necessity of including experimental groups that differed from the initial, neurologically intact cohort across domains (e.g., drug – RX821002 vs. phentolamine, intraperitoneal vs. direct spinal application; preparation – neurologically intact vs. SCI). Given the absence of other relevant sources of data from which to base the power analyses, we used the findings from the neurologically intact, intraperitoneal RX821002 cohort to estimate the sample sizes required for the follow-on cohorts.

For all cohorts, the potential effect of drug on discharge rate was assessed using linear mixed models. The dependent variable for all models was discharge rate. Predictor variables included drug status (pre-vs. post-drug administration; repeated measure), pinch number, electrode channel identifier, and animal. Because multiple pinches were collected for each trial, both before and following drug administration, pinch number was also considered a repeated measure. Random effects included electrode identifier, a drug status by electrode identifier interaction, and a random intercept. Animal was included as a random grouping variable.

We utilized the open-source, web-based General Linear Mixed Model Power and Sample Size (GLIMMPSE) software package(69) to estimate the required number of animals for each cohort. GLIMMPSE enables estimation of sample sizes over a range of target power, main effect, and variability scaling factors. For our analyses, we conservatively assumed that no more than half of the electrodes on each MEA would be considered pinch-responsive. We also assumed only 10 pinches would be collected prior to and following drug administration. We then configured GLIMMPSE to range through target powers of 0.8-1, type 1 error rates of 0.05 and 0.1, scale factors for differences in marginal means of 0.5x, 1x, 1.5x, 2x, corresponding to smaller and larger differences than extracted from neurologically intact rats administered RX821002 intraperitoneally, and variance scale factors of 0.5x, 1x, and 1.5x, also ranging from smaller to larger than expected based on the initial cohort of rats. Sweeping these parameters, the mean estimated sample size across 48 model runs was 7 rats per cohort.

Because each phase of the study introduced a new cohort of animals (except for animals in which the potential impacts both of direct spinal phentolamine and saline administration were assessed), animals were not randomized *a priori.* And because all animals in a given cohort underwent the same experimental protocol, within-cohort randomization was not required. However, this design meant that experimentalists were not fully blind to the drug and/or route of administration for each cohort. However, experimentalists *were* blind to the neural responses to nociceptive sensory feedback being recorded by the MEA, which required subsequent off-line analysis to visualize and quantify. As an additional measure of rigor, one experimentalist performed all pinches for each animal.

This study did not impose animal-level inclusion/exclusion criteria. It did, however, impose electrode channel-level inclusion/exclusion criteria, as described above. Namely, that a given channel must meet the multi-unit voltage and discharge rate metrics required to be considered pinch-responsive. If these qualifications were not met, the channel was not included in the analyses. Single-unit data were sourced from channels already deemed pinch-responsive from the multi-unit records, however discrete neurons from a given channel could be removed from consideration as aforementioned.

For all statistical tests, results were considered significant at the α = 0.05 level. Statistical analyses were performed in SPSS (IBM, Inc.) and GraphPad Prism (GraphPad Software, LLC). Data presented in narrative form are estimated marginal means ± 1 standard error, unless otherwise noted. Graphical depictions of data indicate observed means ± 1 standard deviation, also unless otherwise noted.

## Data availability

Data will be made freely available upon reasonable request.

## AUTHOR CONTRIBUTIONS

JLR: designing research; conducting experiments; acquiring and analyzing data; writing the manuscript. MFB: conducting experiments; acquiring and analyzing data; JGM: designing research; acquiring and analyzing data; writing the manuscript; securing funding.

## ACKNOWLEDGEMENTS

This work was funded by National Institutes of Health grants R01NS111234, R01NS111234-04S1, and R01NS111234-04S2, to J.G.M

## References

1. Llorca-Torralba M, Borges G, Neto F, Mico JA, and Berrocoso E. Noradrenergic Locus Coeruleus pathways in pain modulation. Neuroscience. 2016;338:93–113.

2. Pertovaara A. Noradrenergic pain modulation. Prog Neurobiol. 2006;80(2):53–83.

3. Baker JG, and Summers RJ. Adrenoceptors: Receptors, Ligands and Their Clinical Uses, Molecular Pharmacology and Assays. Handb Exp Pharmacol. 2024;285:55–145.

4. Furst S. Transmitters involved in antinociception in the spinal cord. Brain Res Bull. 1999;48(2):129-41.

5. Kawasaki Y, Kumamoto E, Furue H, and Yoshimura M. Alpha 2 adrenoceptor-mediated presynaptic inhibition of primary afferent glutamatergic transmission in rat substantia gelatinosa neurons. Anesthesiology. 2003;98(3):682–9.

6. Sonohata M, Furue H, Katafuchi T, Yasaka T, Doi A, Kumamoto E, et al. Actions of noradrenaline on substantia gelatinosa neurones in the rat spinal cord revealed by in vivo patch recording. J Physiol. 2004;555(Pt 2):515–26.

7. Pan YZ, Li DP, and Pan HL. Inhibition of glutamatergic synaptic input to spinal lamina II(o) neurons by presynaptic alpha(2)-adrenergic receptors. J Neurophysiol. 2002;87(4):1938–47.

8. Bravo L, Llorca-Torralba M, Berrocoso E, and Mico JA. Monoamines as Drug Targets in Chronic Pain: Focusing on Neuropathic Pain. Front Neurosci. 2019;13:1268.

9. Pertovaara A. The noradrenergic pain regulation system: a potential target for pain therapy. Eur J Pharmacol. 2013;716(1-3):2–7.

10. Starke K, Gothert M, and Kilbinger H. Modulation of neurotransmitter release by presynaptic autoreceptors. Physiol Rev. 1989;69(3):864–989.

11. Mateo Y, and Meana JJ. Determination of the somatodendritic alpha2-adrenoceptor subtype located in rat locus coeruleus that modulates cortical noradrenaline release in vivo. Eur J Pharmacol. 1999;379(1):53–7.

12. Norenberg W, Schoffel E, Szabo B, and Starke K. Subtype determination of soma-dendritic alpha2-autoreceptors in slices of rat locus coeruleus. Naunyn Schmiedebergs Arch Pharmacol. 1997;356(2):159–65.

13. Guyenet PG, Stornetta RL, Riley T, Norton FR, Rosin DL, and Lynch KR. Alpha 2A-adrenergic receptors are present in lower brainstem catecholaminergic and serotonergic neurons innervating spinal cord. Brain Res. 1994;638(1-2):285–94.

14. Rosin DL, Zeng D, Stornetta RL, Norton FR, Riley T, Okusa MD, et al. Immunohistochemical localization of alpha 2A-adrenergic receptors in catecholaminergic and other brainstem neurons in the rat. Neuroscience. 1993;56(1):139–55.

15. Milner TA, Lee A, Aicher SA, and Rosin DL. Hippocampal alpha2a-adrenergic receptors are located predominantly presynaptically but are also found postsynaptically and in selective astrocytes. J Comp Neurol. 1998;395(3):310–27.

16. Fairbanks CA, Stone LS, and Wilcox GL. Pharmacological profiles of alpha 2 adrenergic receptor agonists identified using genetically altered mice and isobolographic analysis. Pharmacol Ther. 2009;123(2):224–38.

17. Reimann W, and Schneider F. Presynaptic alpha 2-adrenoceptors modulate the release of [3H]noradrenaline from rat spinal cord dorsal horn neurones. Eur J Pharmacol. 1989;167(1):161–6.

18. Li X, Zhao Z, Pan HL, Eisenach JC, and Paqueron X. Norepinephrine release from spinal synaptosomes: auto-alpha2 -adrenergic receptor modulation. Anesthesiology. 2000;93(1):164–72.

19. Umeda E, Satoh T, Nagashima H, Potter PE, Tarkovacs G, and Vizi ES. alpha 2A subtype of presynaptic alpha 2-adrenoceptors modulates the release of [3H]-noradrenaline from rat spinal cord. Brain Res Bull. 1997;42(2):129–32.

20. Klimscha W, Tong C, and Eisenach JC. Intrathecal alpha 2-adrenergic agonists stimulate acetylcholine and norepinephrine release from the spinal cord dorsal horn in sheep. An in vivo microdialysis study. Anesthesiology. 1997;87(1):110–6.

21. Olave MJ, and Maxwell DJ. An investigation of neurones that possess the alpha 2C-adrenergic receptor in the rat dorsal horn. Neuroscience. 2002;115(1):31–40.

22. Stone LS, Broberger C, Vulchanova L, Wilcox GL, Hokfelt T, Riedl MS, et al. Differential distribution of alpha2A and alpha2C adrenergic receptor immunoreactivity in the rat spinal cord. J Neurosci. 1998;18(15):5928–37.

23. Howe JR, Yaksh TL, and Tyce GM. Intrathecal 6-hydroxydopamine or cervical spinal hemisection reduces norepinephrine content, but not the density of alpha 2-adrenoceptors, in the cat lumbar spinal enlargement. Neuroscience. 1987;21(2):377–84.

24. Roudet C, Mouchet P, Feuerstein C, and Savasta M. Normal distribution of alpha 2-adrenoceptors in the rat spinal cord and its modification after noradrenergic denervation: a quantitative autoradiographic study. J Neurosci Res. 1994;39(3):319–29.

25. Clarke RW, and Harris J. RX 821002 as a tool for physiological investigation of alpha(2)-adrenoceptors. CNS Drug Rev. 2002;8(2):177–92.

26. Mokha SS, McMillan JA, and Iggo A. Pathways mediating descending control of spinal nociceptive transmission from the nuclei locus coeruleus (LC) and raphe magnus (NRM) in the cat. Exp Brain Res. 1986;61(3):597–606.

27. Tsuruoka M, Maeda M, Nagasawa I, and Inoue T. Spinal pathways mediating coeruleospinal antinociception in the rat. Neurosci Lett. 2004;362(3):236–9.

28. Guo TZ, Jiang JY, Buttermann AE, and Maze M. Dexmedetomidine injection into the locus ceruleus produces antinociception. Anesthesiology. 1996;84(4):873–81.

29. Nicholas AP, Pieribone V, and Hokfelt T. Distributions of mRNAs for alpha-2 adrenergic receptor subtypes in rat brain: an in situ hybridization study. J Comp Neurol. 1993;328(4):575–94.

30. Proudfit HK, and Clark FM. The projections of locus coeruleus neurons to the spinal cord. Prog Brain Res. 1991;88:123–41.

31. Gassner M, Ruscheweyh R, and Sandkuhler J. Direct excitation of spinal GABAergic interneurons by noradrenaline. Pain. 2009;145(1-2):204–10.

32. Gautam M, Yamada A, Yamada AI, Wu Q, Kridsada K, Ling J, et al. Distinct local and global functions of mouse Abeta low-threshold mechanoreceptors in mechanical nociception. Nat Commun. 2024;15(1):2911.

33. Surmeier DJ, Honda CN, and Willis WD, Jr. Natural groupings of primate spinothalamic neurons based on cutaneous stimulation. Physiological and anatomical features. J Neurophysiol. 1988;59(3):833–60.

34. Acuna MA, Kasanetz F, De Luna P, Falkowska M, and Nevian T. Principles of nociceptive coding in the anterior cingulate cortex. Proc Natl Acad Sci U S A. 2023;120(23):e2212394120.

35. Pertovaara A, Hamalainen MM, Kauppila T, Mecke E, and Carlson S. Dissociation of the alpha 2-adrenergic antinociception from sedation following microinjection of medetomidine into the locus coeruleus in rats. Pain. 1994;57(2):207–15.

36. Fung SJ, Manzoni D, Chan JY, Pompeiano O, and Barnes CD. Locus coeruleus control of spinal motor output. Prog Brain Res. 1991;88:395–409.

37. Kauppila T, Jyvasjarvi E, Hamalainen MM, and Pertovaara A. The effect of a selective alpha2-adrenoceptor antagonist on pain behavior of the rat varies, depending on experimental parameters. Pharmacol Biochem Behav. 1998;59(2):477–85.

38. Conway BA, Hultborn H, Kiehn O, and Mintz I. Plateau potentials in alpha-motoneurones induced by intravenous injection of L-dopa and clonidine in the spinal cat. J Physiol. 1988;405:369–84.

39. Rank MM, Murray KC, Stephens MJ, D’Amico J, Gorassini MA, and Bennett DJ. Adrenergic receptors modulate motoneuron excitability, sensory synaptic transmission and muscle spasms after chronic spinal cord injury. J Neurophysiol. 2011;105(1):410–22.

40. Trendelenburg AU, Wahl CA, and Starke K. Antagonists that differentiate between alpha 2A-and alpha 2D-adrenoceptors. Naunyn Schmiedebergs Arch Pharmacol. 1996;353(3):245–9.

41. Convents A, De Keyser J, De Backer JP, and Vauquelin G. [3H]rauwolscine labels alpha 2-adrenoceptors and 5-HT1A receptors in human cerebral cortex. Eur J Pharmacol. 1989;159(3):307–10.

42. Doxey JC, Lane AC, Roach AG, and Virdee NK. Comparison of the alpha-adrenoceptor antagonist profiles of idazoxan (RX 781094), yohimbine, rauwolscine and corynanthine. Naunyn Schmiedebergs Arch Pharmacol. 1984;325(2):136–44.

43. De Vos H, Czerwiec E, De Backer JP, De Potter W, and Vauquelin G. [3H]rauwolscine behaves as an agonist for the 5-HT1A receptors in human frontal cortex membranes. Eur J Pharmacol. 1991;207(1):1–8.

44. Newman-Tancredi A, Nicolas JP, Audinot V, Gavaudan S, Verriele L, Touzard M, et al. Actions of alpha2 adrenoceptor ligands at alpha2A and 5-HT1A receptors: the antagonist, atipamezole, and the agonist, dexmedetomidine, are highly selective for alpha2A adrenoceptors. Naunyn Schmiedebergs Arch Pharmacol. 1998;358(2):197–206.

45. Dennis SG, Melzack R, Gutman S, and Boucher F. Pain modulation by adrenergic agents and morphine as measured by three pain tests. Life Sci. 1980;26(15):1247–59.

46. Kanui TI, Tjolsen A, Lund A, Mjellem-Joly N, and Hole K. Antinociceptive effects of intrathecal administration of alpha-adrenoceptor antagonists and clonidine in the formalin test in the mouse. Neuropharmacology. 1993;32(4):367–71.

47. Cicero TJ, Meyer ER, and Smithloff BR. Alpha adrenergic blocking agents: anti-nociceptive activity and enhancement of morphine-induced analgesia. J Pharmacol Exp Ther. 1974;189(1):72–82.

48. Wei H, and Pertovaara A. Spinal and pontine alpha2-adrenoceptors have opposite effects on pain-related behavior in the neuropathic rat. Eur J Pharmacol. 2006;551(1-3):41–9.

49. Ogilvie J, Simpson DA, and Clarke RW. Tonic adrenergic and serotonergic inhibition of a withdrawal reflex in rabbits subjected to different levels of surgical preparation. Neuroscience. 1999;89(4):1247–58.

50. Choi S, Hachisuka J, Brett MA, Magee AR, Omori Y, Iqbal NU, et al. Parallel ascending spinal pathways for affective touch and pain. Nature. 2020;587(7833):258-63.

51. Dhandapani R, Arokiaraj CM, Taberner FJ, Pacifico P, Raja S, Nocchi L, et al. Control of mechanical pain hypersensitivity in mice through ligand-targeted photoablation of TrkB-positive sensory neurons. Nat Commun. 2018;9(1):1640.

52. Lin Q, Peng YB, and Willis WD. Antinociception and inhibition from the periaqueductal gray are mediated in part by spinal 5-hydroxytryptamine(1A) receptors. J Pharmacol Exp Ther. 1996;276(3):958–67.

53. Nadeson R, and Goodchild CS. Antinociceptive role of 5-HT1A receptors in rat spinal cord. Br J Anaesth. 2002;88(5):679–84.

54. Gjerstad J, Tjolsen A, and Hole K. The effect of 5-HT1A receptor stimulation on nociceptive dorsal horn neurones in rats. Eur J Pharmacol. 1996;318(2-3):315–21.

55. You HJ, Colpaert FC, and Arendt-Nielsen L. The novel analgesic and high-efficacy 5-HT1A receptor agonist F 13640 inhibits nociceptive responses, wind-up, and after-discharges in spinal neurons and withdrawal reflexes. Exp Neurol. 2005;191(1):174–83.

56. Besnard J, Ruda GF, Setola V, Abecassis K, Rodriguiz RM, Huang XP, et al. Automated design of ligands to polypharmacological profiles. Nature. 2012;492(7428):215-20.

57. Milne B, Sutak M, Cahill CM, and Jhamandas K. Low dose alpha-2 antagonist paradoxically enhances rat norepinephrine and clonidine analgesia. Anesth Analg. 2011;112(6):1500–3.

58. Lilius TO, Rauhala PV, Kambur O, Rossi SM, Vaananen AJ, and Kalso EA. Intrathecal atipamezole augments the antinociceptive effect of morphine in rats. Anesth Analg. 2012;114(6):1353–8.

59. Gordh T, Jr., Jansson I, Hartvig P, Gillberg PG, and Post C. Interactions between noradrenergic and cholinergic mechanisms involved in spinal nociceptive processing. Acta Anaesthesiol Scand. 1989;33(1):39–47.

60. Xu Z, Li P, Tong C, Figueroa J, Tobin JR, and Eisenach JC. Location and characteristics of nitric oxide synthase in sheep spinal cord and its interaction with alpha(2)-adrenergic and cholinergic antinociception. Anesthesiology. 1996;84(4):890–9.

61. Levine JD, Taiwo YO, Collins SD, and Tam JK. Noradrenaline hyperalgesia is mediated through interaction with sympathetic postganglionic neurone terminals rather than activation of primary afferent nociceptors. Nature. 1986;323(6084):158-60.

62. Ogon I, Takebayashi T, Miyakawa T, Iwase T, Tanimoto K, Terashima Y, et al. Attenuation of pain behaviour by local administration of alpha-2 adrenoceptor antagonists to dorsal root ganglia in a rat radiculopathy model. Eur J Pain. 2016;20(5):790–9.

63. Tanimoto K, Takebayashi T, Kobayashi T, Tohse N, and Yamashita T. Does norepinephrine influence pain behavior mediated by dorsal root ganglia?: a pilot study. Clin Orthop Relat Res. 2011;469(9):2568–76.

64. Bandres MF, Gomes J, and McPherson JG. Spontaneous Multimodal Neural Transmission Suggests That Adult Spinal Networks Maintain an Intrinsic State of Readiness to Execute Sensorimotor Behaviors. J Neurosci. 2021;41(38):7978–90.

65. Bandres MF, Gomes JL, and McPherson JG. Spinal stimulation for motor rehabilitation immediately modulates nociceptive transmission. J Neural Eng. 2022;19(5).

66. Bandres MF, Gomes JL, and McPherson JG. Intraspinal microstimulation of the ventral horn has therapeutically relevant cross-modal effects on nociception. Brain Commun. 2024;6(5):fcae280.

67. Quiroga RQ, Nadasdy Z, and Ben-Shaul Y. Unsupervised spike detection and sorting with wavelets and superparamagnetic clustering. Neural Comput. 2004;16(8):1661–87.

68. McPherson JG, and Bandres MF. Spontaneous neural synchrony links intrinsic spinal sensory and motor networks during unconsciousness. Elife. 2021;10.

69. Kreidler SM, Muller KE, Grunwald GK, Ringham BM, Coker-Dukowitz ZT, Sakhadeo UR, et al. GLIMMPSE: Online Power Computation for Linear Models with and without a Baseline Covariate. J Stat Softw. 2013;54(10).

